# Unilateral optogenetic inhibition and excitation of basal ganglia output show opposing effects on directional lick choices and movement initiation in mice

**DOI:** 10.1101/360818

**Authors:** Arthur E. Morrissette, Po-Han Chen, Conrad Bhamani, Peter Y. Borden, Christian Waiblinger, Garrett B. Stanley, Dieter Jaeger

**Affiliations:** Neuroscience Graduate Program, Atlanta, Georgia 30322; Department of Biology, Emory University, Atlanta, Georgia 30322; Department of Biomedical Engineering, Georgia Institute of Technology, Atlanta, Georgia 30332

## Abstract

Models of basal ganglia function predict that tonic inhibitory output to motor thalamus suppresses unwanted movements, and that a decrease in such activity leads to action selection. A direct test of these outcomes of thalamic inhibition has not been performed, however. To conduct such a direct test, we utilized rapid optogenetic activation and inactivation of the GABAergic output of the substantia nigra pars reticulata (SNr) to motor thalamus in mice that were trained in a sensory cued left/right licking task. Directional licking tasks have previously been shown to depend on a thalamocortical feedback loop between ventromedial motor thalamus and antero-lateral premotor cortex (Li et al., 2015; Guo et al., 2017). In confirmation of model predictions, we found that 1s of unilateral optogenetic inhibition of GABAergic output from the SNr biased decision making towards the contralateral lick spout with ipsilaterally cued trials while leaving motor performance intact. In contrast, 1s of optogenetic excitation of SNr terminals in motor thalamus resulted in an opposite bias towards the ipsilateral direction confirming a bidirectional effect of tonic nigral output on directional decision making. In a second variant of the task we disallowed anticipatory licking and found that successful suppression of anticipatory licking was also impacted by our optogenetic manipulations in agreement with the suppressive effect of tonic nigral output. Nevertheless, direct unilateral excitation of SNr cell bodies resulted in bilateral movement suppression, suggesting that descending motor pathways from the SNr to superior colliculus also play an important role in the control of licking behavior.

**Significance Statement:** This study provides the first evidence that basal ganglia output to motor thalamus can control decision making in left/right licking choices and suppress anticipatory movement initiation. Unilateral optogenetic inhibition or excitation of basal ganglia output via the substantia nigra resulted in opposite changes of directional lick choices and could override the sensory information on lick direction provided by a whisker stimulus. These results suggest that basal ganglia output gates activity in a thalamo-cortical feedback loop previously shown to underlie the control of forced choice directional licking behavior. The results substantiate models stating that tonic inhibition of motor thalamus from the basal ganglia directs action selection and suppresses unwanted movements.

## Introduction

The basal ganglia (BG) are thought to influence the control of movement by inhibiting and facilitating selected motor programs to accomplish and reinforce goal-directed behavior (Albin et al., 1989; DeLong, 1990). Our long-standing understanding of the BG’s role in the control of movement suggests that the output nuclei of the BG provide tonic inhibition over motor areas of the thalamus and brainstem to suppress unwanted movements; pauses in the inhibitory output would therefore facilitate movement initiation (Deniau and Chevalier, 1985; Mink, 1996; Hikosaka, 2007). In support of this general basal ganglia functional model, optogenetic activation of the basal ganglia direct pathway has been found to be pro-kinetic, leading to a reduction in the inhibition of the SNr (Freeze et al., 2013), the primary basal ganglia output nucleus in the rodent, and disinhibition of downstream targets (Oldenburg and Sabatini, 2015). Additionally, pauses in SNr activity preceded initiation of orienting movements in rats performing an auditory-cued Go-Stop task (Schmidt et al., 2013). It remains unclear, however, how cortical planning activity and motor initiation is affected by output from the basal ganglia.

Recent studies have shown that in mice directional (left vs. right) licking in delayed choice tasks is initiated from an anterolateral premotor area named ALM (Guo et al., 2014a; Li et al., 2015; Chen et al., 2017). ALM expresses ramping activity in a feedback loop with the ventromedial thalamus (VM) during preparation of directional licking, and inhibition of either area reduces task performance to chance (Guo et al., 2017). The VM thalamus is a major part of the motor thalamus receiving strong input from the basal ganglia (BGMT, which also includes a portion of the anterior VA/VL nuclei) (Kuramoto et al., 2011). The basal ganglia are thus well positioned to control this corticothalamic feedback loop through their strong inhibitory projection to motor thalamus from the substantia nigra reticulata (SNr) and the globus pallidus internus (GPi). Directional licking behavior as planned and initiated by ALM presents therefore an ideal model system to study how the basal ganglia exert motor effects in cortically initiated movements. To address this question, we trained mice to perform a licking task where they must lick a left or right positioned lick spout following a left or right air puff, respectively. In order to further parse the contributions of BG output on cognitive control of movement preparation and suppression, we trained the same mice to perform two variations of this task: first allowing mice to lick before the response window and anticipate the reward delivery (‘anticipatory’ task variant) and later a variant that required mice to actively withhold licking until the onset of the response window (‘withholding’ task variant). By training the mice to perform these task variants one after another, we were able to parse the basal ganglia contributions to initiating or suppressing anticipatory movements and the control of rewarded choice preparation and initiation. Using optogenetic methods to inhibit SNr neurons unilaterally, we found that silencing the inhibitory output of the basal ganglia triggered licking contralateral to the side of inhibition, irrespective of the rewarded licking direction. In contrast, exciting SNr projections to the motor thalamus and inhibiting thalamic activity suppressed contralateral licking and biased licking towards ipsilateral direction. These results suggest that the basal ganglia output via SNr to the motor thalamus exerts powerful unilateral control over movement preparation and initiation in the context of sensory guided motor behavior.

## Methods

### Animals

For optogenetic stimulation and electrophysiological experiments, male and female Vgat-IRES-Cre mice (*Slc32a1*) aged 6-12 months at the start of experiments were used. Mice were maintained on a 12h:12h reverse light cycle and all experiments and behavioral training was performed during the dark portion of the cycle. Mice undergoing behavioral training were provided *ad libitum* food access and were kept on 1-1.5 ml/day water restriction 6 days a week starting at least 3 days prior and for the duration of handling, training, and experimental testing. On day 7 of each week, mice were given free access to water. During behavioral training and testing, mice were given 10% sucrose solution with 0.1% grape Kool-Aid powder. Liquid consumption was measured during testing and mice were supplemented with water to reach the 1-1.5 ml/day volume. All experimental procedures were approved by the Emory University Institutional Animal Care and Use Committee.

### Surgery

#### Viral Vector Injection and Fiber Placement

For optogenetic AAV vector injection, mice were anesthetized with isoflurane (induction at 3-4%, maintained at 1-2%) and head-fixed on a stereotaxic frame (Kopf). Craniotomies were made unilaterally above the SNr. For somatic SNr inhibition, 200nL of AAV5-EF1a-DIO-eARCH3.0-eYFP (ARCH; n = 5, 3 male) or for excitation, 200nL of AAV5-EF1a-DIO-hChR2(E123T/T159C-EYFP (ChR2(ET/TC) or ChR2; n = 4, 2 male) was injected with a nano-injector (Nanoject II, Drummond Scientific) at the rate of 0.46nl/s into the SNr targeting the coordinates (in mm from Bregma): AP −3.2, ML 1.6, DV −4.2). For somatic SNr excitation and SNr-BGMT terminal excitation 200µm, 0.22NA optic fibers (Thor Labs) each with a 1.25mm steel ferrule were then implanted targeting the SNr and the ventromedial thalamus (AP −1.5, ML 0.9, DV −4.0). Following surgical implantation, mice were kept in single housed cages. All optogenetic manipulations were performed unilaterally on the right side of the animal, and hence ‘ipsilateral’ (IPSI) in this paper always refers to the right and ‘contralateral’ (CONTRA) refers to the left side of the body.

#### Acute Electrode/Optrode Recordings

To record and/or optogenetically manipulate neural populations in the right SNr or downstream projection targets, a craniotomy was made above the site targeting these areas and the dura was kept intact. A 4mm diameter and 1mm tall plastic tube was glued in place around the craniotomy and the area was filled with a removable elastomer (Kwik-Cast, World Precision Instruments) to allow for access to the tissue for future experiments.

### Behavior

Prior to behavioral training, a custom stainless-steel head-post was attached posterior to lambda by placing a thin layer of cyanoacrylate adhesive on the skull, followed by a thin layer of dental acrylic (Metabond; C&B Associates). Following recovery, mice were head-fixed and placed within the behavioral training setup consisting of two 2mm diameter lick spouts placed 5mm apart and two 2mm diameter air-puff tubes that were directed at the C row of whiskers. The lick spouts were connected to a custom capacitive lick sensor circuit that recorded time and duration of licks at 200 Hz. Air-puff intensities were calibrated such that whisker deflection was apparent under high-speed video monitoring without signs of freezing or startle behavior from the mouse.

### Behavioral Task

Animal training protocols and behavioral paradigm are adapted from previously reported procedures (Guo et al., 2014b). The behavioral paradigm is illustrated in Figure 1A. Left and right air-puff/lick trials were selected pseudo-randomly such that the probability of a right/left trial was adjusted based on the ratio of left/right rewarded trials (e.g. for the trial history of the current session consisting of 8 correct right trials and 4 correct left trials, the probability that the next trial would be left was set to right rewards/total rewards = 66%). Each trial was made up of 3 discrete intervals: a “pre-stimulus” period, a “sensory” period where mice received either a left or right air-puff for 1-1.5 s, followed by a 5 s “response” period where the mice lick the left or right spout to indicate which spout they believe will be rewarded and then consume the reward if the correct choice was made. Trials were separated by a variable inter-trial period. During the sensory period, mild non-aversive air-puff stimuli were directed through 2mm-diameter steel tubes towards the whiskers. The tubes were angled at 15 degrees away from the mouse center to isolate air-puff stimuli to the whiskers, avoiding the face of the mouse. The stimuli lasted for 1-1.5 s during training, though for optogenetic experiments during the withholding task, the sample period was held constant at 1 s.

**Figure 1.**
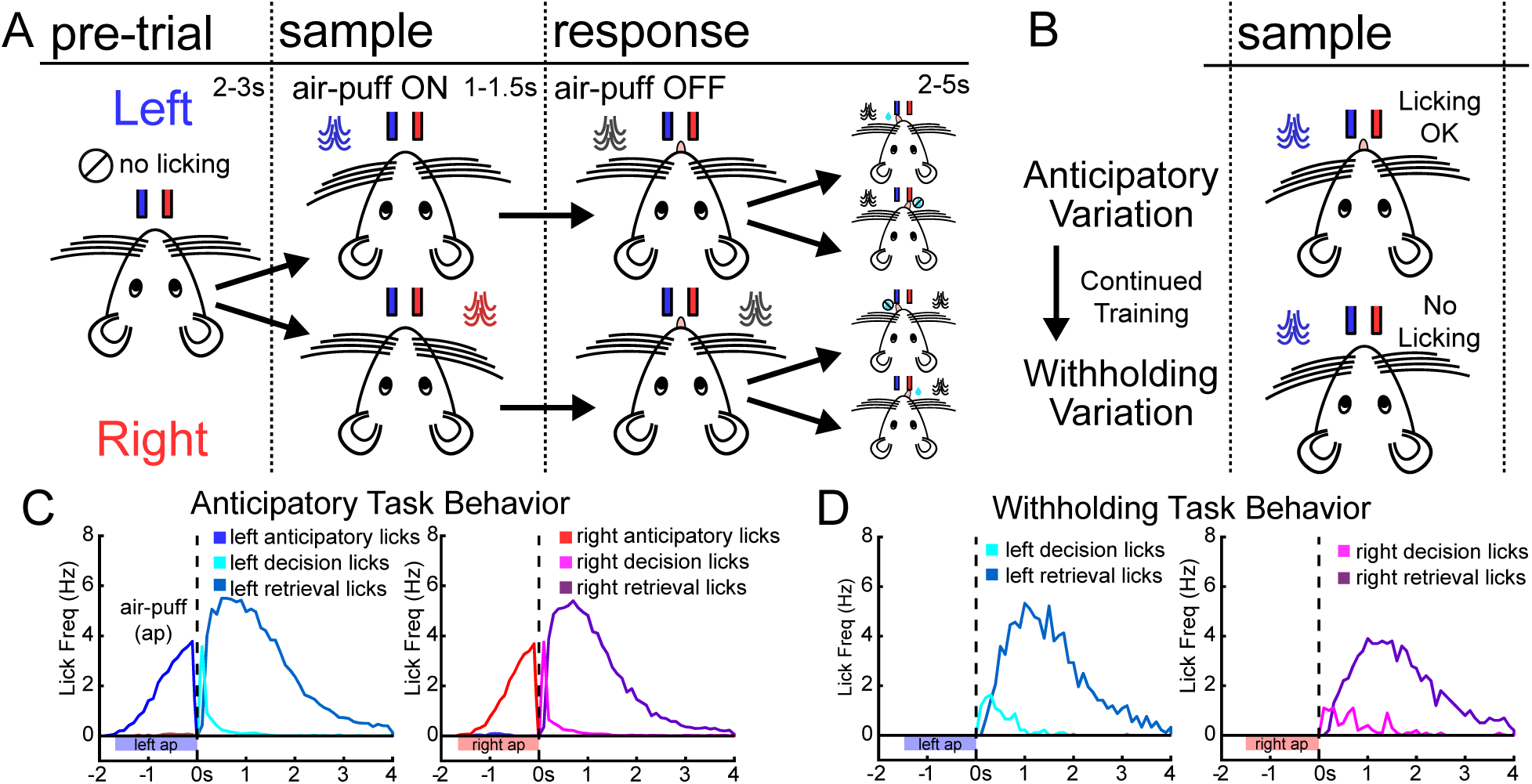
Mice learn to perform anticipatory and withholding variants of a forced choice licking task. **A.** Illustration of the forced choice licking task. Each trial begins with a pre-trial period lasting 2-3 s where the mouse cannot lick either spout. During the sample period, an air-puff is directed at either the left or right whiskers and remains on for 1-1.5 s. The response period begins after the offset of the air-puff and the first lick during this period is counted as the decision lick. If the decision lick is correct (towards the left spout for left air-puff or right lick spout for right air-puff) the mouse is provided a liquid reward from the correct spout. If the mouse does not provide a lick for the duration of the response period (2-5 s) or licks the incorrect spout no reward is provided. **B.** Mice are trained in two subsequent variations of the licking task. First, they are trained in the anticipatory variation where they are allowed to lick towards either spout during the air-puff in anticipation of the response period and potential reward delivery. After they have learned this task, the same mice are further trained in the withholding variation of the task where the must suppress licking during the duration of the air-puff. **C.** Average lick frequency traces for correct trials from mice performing the anticipatory variant of the task (N = 12 mice). **Left.** For left trials (n = 1151 trials), Mice begin to show anticipatory licking (middle blue) after the onset of the air-puff and the first lick after the offset of the air-puff is counted as the decision lick (light blue). The retrieval licks (dark blue) are the licks corresponding to the mouse retrieving the sucrose reward after a correct decision lick. **Right.** Same format as left trials, but instead showing right trials (n = 1098). Anticipatory, decision, and retrieval licks are depicted as middle red, light red, and dark red, respectively. **D.** Average lick frequency traces for correct trials from mice performing the withholding variant of the task (N = 8 mice). **Left.** For left trials (n = 397 trials), mice successfully withhold licking activity during the air-puff and once again the first lick after the offset of the air-puff is counted as the decision lick (light blue). The retrieval licks (dark blue) are the licks corresponding to the mouse retrieving the sucrose reward after a correct decision lick. **Right.** Same as for left, but now for right lick trials (n = 403 trials).

Training for the behavioral task began after a minimum of 10 days following surgery and 5 days following the start of water restriction. Mice were handled during initial days of water restriction to acclimate with handling. On the first day of head-fixation and training, mice were secured in the holder and placed in the behavioral setup with the two lick spouts positioned and centered in front of the mouth of the mouse. During the first day of training, mice could lick both the left or right spout and would receive a sucrose reward following licks at a minimum interval of 10 s. This was primarily used to acclimate the mouse with head-fixation and the positions of the lick spouts. The positions of the spouts were manually adjusted relative to the center of the mouth to promote equal licking of both lick spouts. During subsequent days of training, mice were transitioned to a task where the air-puffs signifying the rewarded spout were active, though the reward was automatically triggered (“Auto-Reward”). This typically lasted 1 day of training (∼4 50 trial blocks) until the mice became acclimated to the puffs and would display anticipatory licking towards the correct lick spout before the reward was triggered, demonstrating that the mice had built the association between air-puff location and rewarded lick spout. In the next step of training, typically occurring on days 2-3, mice received the air-puffs and were then allowed to lick freely towards both spouts in the Response period and were only rewarded on licks to the correct spout (“Free-Lick”). Mice during this time learned to make the correct perceptive decision (signified by licking the rewarded spout exclusively during trials). Mice were then introduced to the ‘anticipatory’ variant of the task (Fig 1C). This variant of the task now penalized incorrect decision licks during the reward period by transitioning straight into a lengthened inter-trial delay period (6-10 s vs. 2-4 s for passed trials). To minimize licking outside of the response interval of the trials, each trial began with a pre-stimulus period where the mouse was required to withhold licking for at least 2-3 s before air-puff onset. Licking during the last second of this period led to a trial fail and mice were placed back in the inter-trial period awaiting the start of the next trial. Training mice to perform the anticipatory task occurred over 2-5 days. Finally, mice were trained on the second variant of the task that required the mice to withhold licking (‘withholding’ variant) during the air-puff stimulus period until the offset of the air-puff (Fig 1D). To train mice to begin withholding licks during the air-puff stimulus period, any lick activity during the air-puff triggered an “early lick” fail and was penalized with an extended inter-trial period. At the beginning, the air-puff was shortened to 0.5 s and gradually increased in steps of 0.25 s until the mice were able to withhold licking and achieve 60% performance with a variable sample period duration between 1-1.5 s. This training typically took an additional 1-2 weeks. Of the 12 mice that were trained in the anticipatory variant of the task, 8 mice were able to subsequently learn the withholding variant.

### Optogenetic stimulation

Before and after each session the output intensity of the light source (either LED or laser) was determined using an optical power meter and sensor (PM100D and S121C, ThorLabs). For SNr inhibition experiments, we used a 593nm yellow laser (Shanghai Dream Lasers) collimated and coupled to a 200micron, 0.22 NA patch cable (Doric Lenses) leading to 8-12mW output from the fiber tip. For somatic SNr excitation we used a 470nm LED (Doric Lenses) coupled to the same patch cable with output power between 1-3mW from fiber tip. Finally, for SNr-BGMT terminal excitation, we used a 470nm blue laser (Shanghai Dream Lasers) with the same patch cables with output power between 8-12mW from the fiber tip. Trials with optogenetic stimulation were randomly intermixed with control trials for a total proportion between 25-50% of trials with light exposure. The optical stimulation trials were also all executed with a fixed 1 s air-puff duration in order to avoid inconsistent relations between stimulation and the timing of anticipatory licking.

### Electrophysiology

During surgical preparation, a craniotomy was made over the future right SNr/BGMT recording sites (−4 to 1mm AP, 0.5 to 2.5mm ML) and covered with Kwik-cast (WPI Inc.). The dura was left intact. A stainless-steel reference skull screw (#19010-10, Fine Science Tools) was placed over the contralateral sensory cortex. A 0.01” diameter steel wire was soldered between the screw and a gold pin to connect to the acquisition system during recording. The mice were allowed at least 3 days to recover. Following recovery, mice were acclimated to being head-fixed and recording sessions began (one session per day, 2 hours per session). Within the first session of head-fixation, mice showed no overt signs of stress and appeared relaxed. During recording, mice were maintained on a randomized interval reward paradigm where mice were provided with a sucrose reward via right or left lick spout every 30-60 s to encourage quiet wakefulness during the session. At the start of the session, the Kwik-cast cover over the craniotomy was removed. Custom optrodes consisting of a 50-100 micron optic fiber (ThorLabs) attached 200-300 microns above a micro-electrode (FHC) were lowered into the SNr or BGMT. Raw signals (0.1–10 kHz band-pass filtered) were acquired at 20 kHz, amplified and digitized (RHD2132 headstage, Intan Technologies) and saved (RHD2000 Evaluation System/Interface Software, Intan Technologies). Once unit activity was detected in the SNr or BGMT, optical stimulation with either a yellow (593 nm) or blue (473 nm) laser was delivered for 1 or 2 s continuous pulses every 10 s to stimulate ARCH3 or ChR2 expressing neurons, respectively. For some recordings, the optic fiber was placed in the SNr with a separate electrode lowered in to the BGMT for recording activity downstream of the site of optogenetic stimulation. After each session, the craniotomy was covered with Kwik-cast and following the final recording session, the mouse was perfused with PBS followed by perfusion with 4% paraformaldehyde and 15% sucrose. The brain was then removed and transferred to a 4% paraformaldehyde/30% sucrose solution for later histological processing.

### Data Analysis

Analysis of behavioral and electrophysiological data was performed in MATLAB (MathWorks). Only behavioral sessions where baseline performance was above 60% were included in analysis. Behavioral trials in which the mouse licked <1 s before the start of the trial (onset of the air-puff) were caught and sent to inter-trial delay. These trials were rare (<2% in trained mice and excluded from analysis). Lick data was pre-processed to remove “artifact licks” (spout contacts shorter than 10 ms and contacts lasting longer than 200 ms typically caused by electrical noise and paw touches, respectively). Trials where the decision lick (first lick during the response period) was classified as an artifact lick were removed from subsequent analyses. To calculate average lick frequencies for the various behavioral and experimental conditions, the onsets of lick contacts were marked and lick contacts across each trial were summed in 50 ms bins and divided by the length of the bin duration. Significance of the performance change in each optogenetic stimulation condition was determined using bootstrapping to account for variability across mice, sessions, and trials. We tested against the null hypothesis that the performance change caused by optogenetic stimulation was due to normal behavioral variability. In each round of bootstrapping, we replaced the original data set with a re-sampled set in which we resampled with replacement from: 1) animals; 2) sessions performed by each animal; and 3) the trials within each session. We then computed the change in performance on the re-sampled data set. Repeating this procedure 10,000 times produced a distribution of performance changes that reflected the behavioral variability. The P value observed performance change was calculated as the fraction of times bootstrapping produced an inconsistent performance change (for example, if a performance decrease was observed during optogenetic stimulation, the P value is the fraction of times a performance increase was observed during bootstrapping, one-tailed test). Error bars represent the +/-SEM generated from bootstrapping unless noted otherwise.

## Results

### Mice learn to perform ‘anticipatory’ and ‘withholding’ variants of a forced choice licking task

In order to understand how the basal ganglia output influences directed licking behavior, we employed a forced choice behavioral task. In order to further parse the contributions of BG output on cognitive control of movement preparation and suppression, we trained the same groups of mice to perform two variations of this task: first, allowing mice to lick before the response window and anticipate the reward delivery, and second, a new variant that required mice to actively withhold licking until the onset of the response period (Fig 1B). By training the same mice to perform these task variations in sequence, we were able to parse the basal ganglia contributions to anticipation and initiation of licking movements (anticipatory task) as well as the control of movement preparation and suppression (withholding task).

### Unilateral inactivation of the SNr biases towards contralateral licking behavior

We began assessing the role of basal ganglia output in the anticipatory variant of the forced choice licking task using fast, reversible optogenetic inactivation of the SNr through nigral ARCH3 activation (Fig 2A). Cre-dependent ARCH3 was injected to the SNr of VGAT-cre mice that express cre in GABAergic populations. Extent of expression was well localized to the SNr with axon terminals extending towards known projection targets of the SNr such as the substantia nigra compacta, brainstem and thalamus (typical example shown in Fig 2A, right). In awake mice at rest, yellow (593nm) light stimulation of SNr neurons expressing ARCH3 abolished nearly all firing activity for the duration of the stimulation (mean firing rates 17.2Hz baseline vs. 2.61Hz with ARCH inhibition, n = 10 single units, Fig 2BC). Following the offset of the optogenetic stimulation, the firing rate quickly returned to baseline levels. In behaving mice, we optogenetically manipulated the SNr unilaterally in the right hemisphere (Fig 2D). To examine the role of the SNr in anticipatory licking activity, the SNr was inhibited from 0.5 s before the offset of the air-puff until 0.5 s into the response period of the ‘anticipatory’ variant of the task. Inhibiting the right SNr during this period biased licking activity towards the contralateral direction (Fig 2E,H). A striking increase in ipsilateral air-puff trials with contralateral decision licks was observed and the overall success rate of ipsilateral trials was decreased (Fig 2H). Additionally, the decrease in successful trial performance with ipsilateral air-puffs was due to an increase in no-response trials (Fig 2H, P<0.001). In contrast, contralateral air-puff trials showed an increase in successes associated with a decrease in mistaken decision licks to the ipsilateral direction (5 mice, 36 sessions, P<0.001 (left), P<0.001 (right), bootstrap, Fig 2H). Importantly, successful trials showed the same amount and timing of anticipatory licking and the same consummatory lick rate in trials with air-puffs on either side with or without SNr inhibition (Fig 2F). This suggests that the lateralization of decision making towards the spout contralateral to SNr inhibition was not due to a slowing of movement, but rather a categorical change in the decision process. In control mice expressing injected with an EYFP virus without ARCH3, light stimulation had no effect on licking behavior or task performance (2 mice, 11 sessions, data not shown).

**Figure 2.**
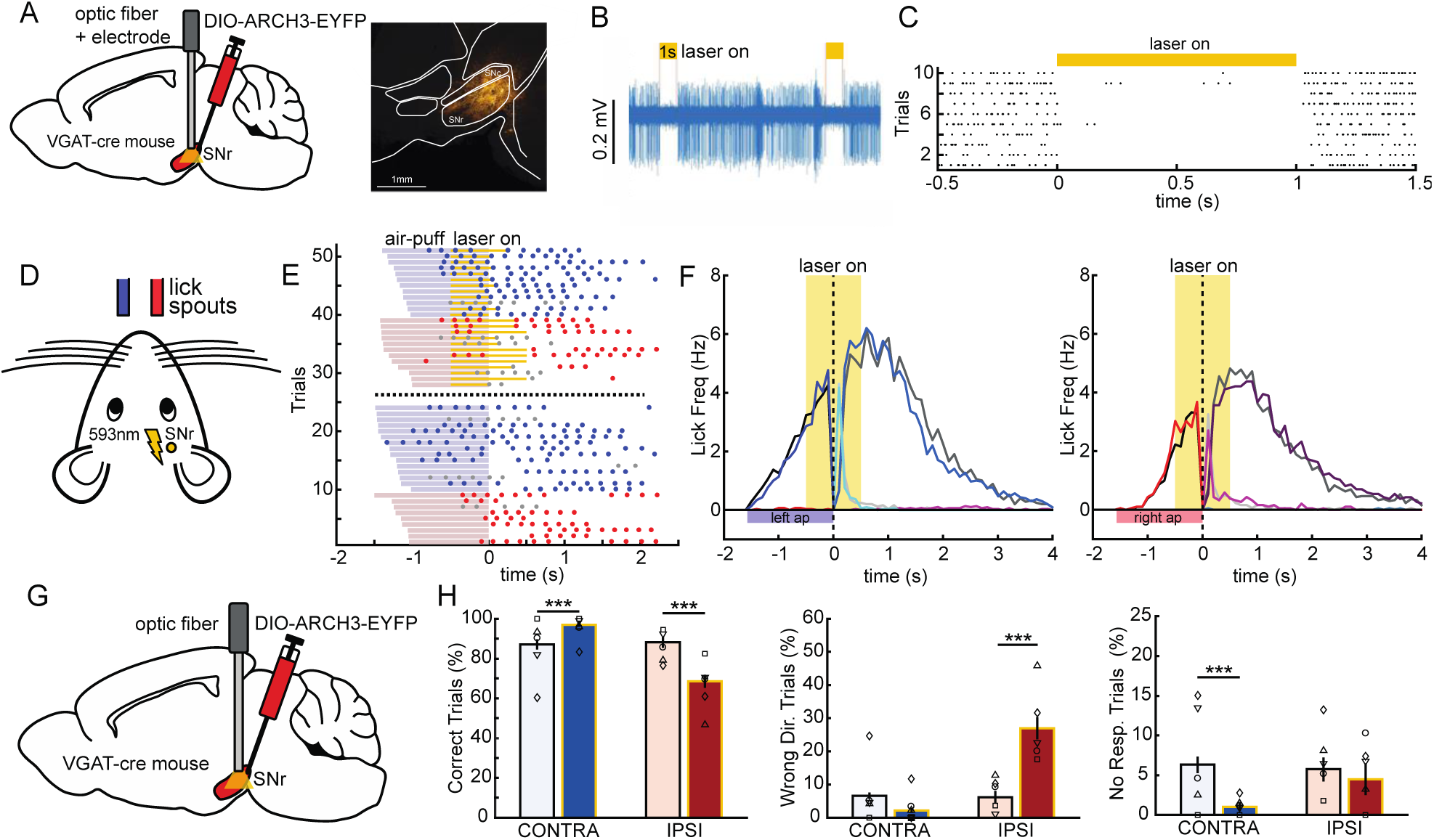
Unilateral inactivation of the SNr biases towards contralateral licking behavior. **A.** Diagram of optogenetic vector injection and optic fiber targeting. Cre-dependent AAV2-DIO-ARCH3-EYFP was injected in the right SNr of VGAT-cre transgenic mice and optic fiber attached to a microelectrode was lowered into SNr. Example of ARCH3-EYFP (yellow) expression in SNr (right). **B.** Example single-unit recording of an SNr neuron expressing ARCH3. 593nm laser stimulation silences firing activity for the duration of the light pulse and firing quickly resumes following the offset of the light stimulation. **C.** 10 stimulation trials aligned to the onset of stimulation that show the consistency of ARCH3 inhibition of SNr activity. **D.** Illustration showing mouse orientation with respect to ipsilateral (red) and contralateral (blue) lick spouts positioned in front of the mouse and optogenetic illumination through fiber implanted over the SNr. **E.** Example behavioral session showing licking activity for ipsilateral and contralateral optogenetic and baseline trials. Trials are arranged by optogenetic stimulation condition (stim trials top, baseline below) and length of air-puff presentation. Blue and red horizontal lines depict the presentation of the air-puff during the sample period and yellow lines are the periods of optogenetic SNr inhibition. In this example, optogenetic inhibition of right SNr disrupts right (ipsilateral) licking behavior such that the mouse is biased towards licking the contralateral spout. **F.** Average lick frequency traces for correct trials with (colored lines) and without (gray lines) optogenetic SNr inhibition. For contralateral trials (n = 300 opto, 247 baseline, **left**) and ipsilateral trials (n = 185 opto, 301 baseline, **right**) mice show anticipatory licking activity towards the correct spout that increases until the start of the response period. Anticipatory licks are depicted by the middle colored line, followed by decision licks (light color) and retrieval licks (dark colored line). **G.** Behavioral effects of unilateral SNr optogenetic inhibition in the anticipatory task. **H.** Changes in performance between contralateral (blue) and ipsilateral (red), off (light) and on (dark) stimulation. Bar height represents mean across all sessions (n = 35 sessions), with shapes representing the mean for each mouse (5 mice). Error bars represent SEM (bootstrap, 10000 iterations) and p values based on bootstrap (see methods, ***p<0.001, **p<0.01, *p<0.05). **Left.** B Compared to interleaved baseline trials (light blue/red columns), licking performance during unilateral SNr inhibition (dark blue/red columns) produced a significant increase in the percent of contralateral trials correct (p<0.001) and significant decrease in the percent of ipsilateral trials correct (35 sessions, 5 mice). **Middle.** Optogenetic inhibition of the SNr caused a significant increase (P<0.001) in the percent of ipsilateral wrong direction fail trials (incorrectly lick contralateral spout during ipsilateral trials). **Right.** For contralateral lick trials, SNr inhibition significantly reduced the number of no response trials (P<0.001).

### Unilateral inhibition of the SNr increases anticipatory licks and decreases reaction time towards the contralateral direction

To test the hypothesis that SNr optogenetic inhibition interferes with the ability to withhold unwanted anticipatory movements, we activated ARCH during the 2^nd^half of the air-puff stimulus in the same mice after further training them to perform the ‘withholding’ variant of the task. A subset (3 of 5) of the mice were able to learn this variant of the task. After achieving 60% performance, we again tested the effect of optogenetic manipulation starting 0.5 s before the offset of the 1 s air-puff. This manipulation again lateralized the execution of trials, but with additional features compared to the ‘anticipatory’ task (Fig 3). Ipsilateral air-puff trials again resulted in a significantly increased proportion of wrong direction decision licks (Fig 3E, P<0.01, bootstrap). Nevertheless, overall correct performance in the withholding version of the task was slightly increased in ipsilateral air-puff trials with optogenetic SNr inhibition (3 mice, 17 sessions, P=0.041, bootstrap), opposite of the observed change in the anticipatory task. By further examining the breakdown of failure types (Fig 3E), we found a reduction of failures due to early licking (P<0.001, bootstrap), indicating a higher success rate of suppressing anticipatory licking in the ‘withholding’ task when SNr was inhibited. In contrast, contralateral air-puff trials were associated with a significant increase in such early licks (P<0.01, bootstrap). This supports the hypothesis that inhibiting SNr output did not only have direct consequences for on the decision of lick direction, but also impaired the ability to withhold licks towards the direction contralateral to optogenetic stimulation and facilitated withholding to the ipsilateral direction. The average lick frequency traces from correct trials (Fig 3C) with optogenetic stimulation were again similar to control trials, suggesting that the licking process when initiated was not itself disrupted. However, success trials did show a shorter reaction time for contralateral licks and a longer reaction time for ipsilateral licks compared to baseline trials (Fig 3 FG, P<0.01 (left), P<0.01 (right), bootstrap). This suggests that when the mice were able to withhold licking during the optogenetic stimulation, suppressing basal ganglia output biases preparation of movement or motor readiness, such that the mouse was able to respond more quickly to the contralateral spout and more slowly towards the ipsilateral direction. Control mice expressing EYFP in the SNr but no ARCH (n = 2 mice, data not shown), showed no significant changes in trial outcomes between baseline and optogenetic stimulation trials.

**Figure 3.**
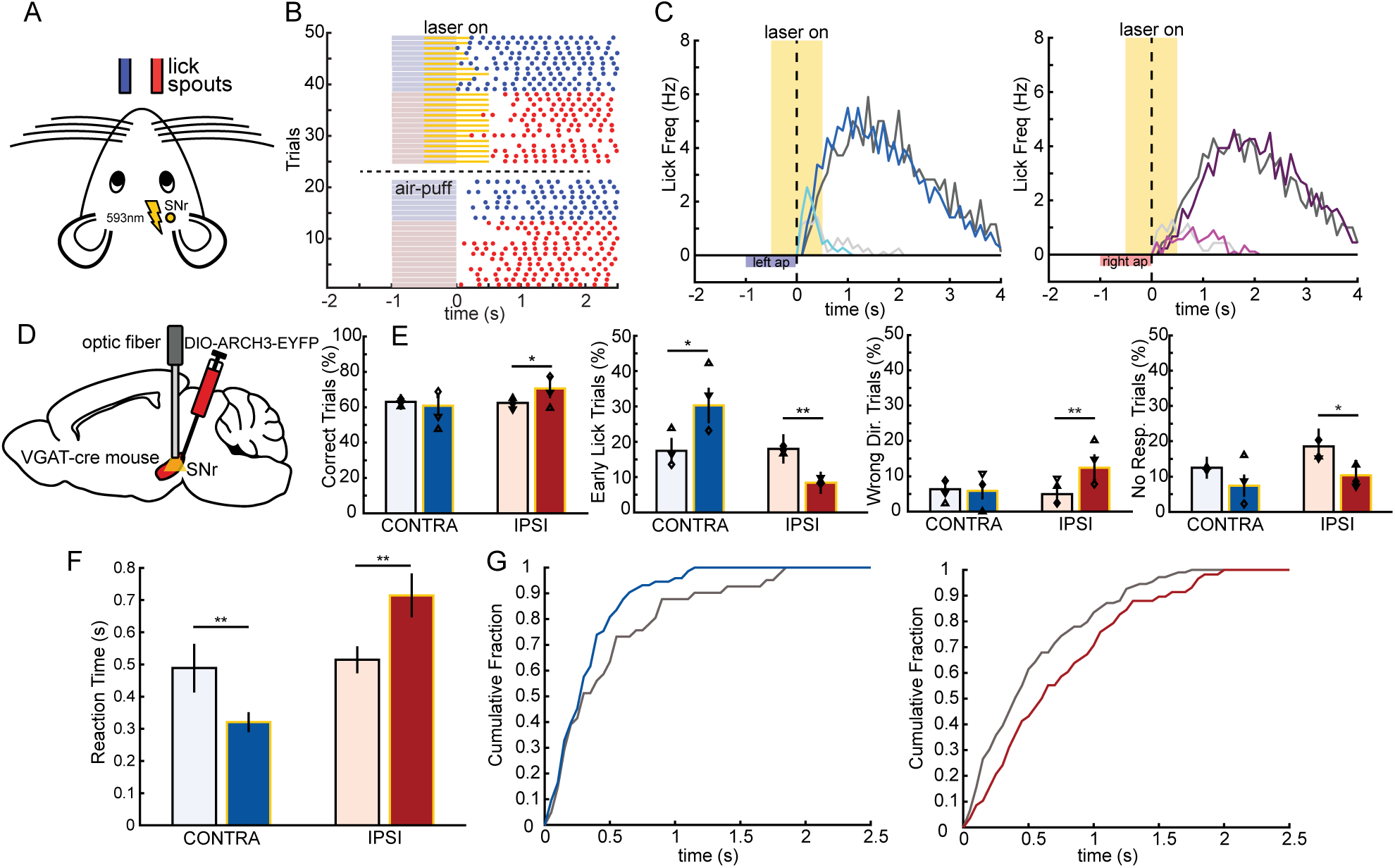
Unilateral inhibition of the SNr increases anticipatory licks and decreases reaction time towards the contralateral direction. **A.** Illustration showing mouse orientation with respect to ipsilateral (red) and contralateral (blue) lick spouts positioned in front of the mouse and optogenetic illumination through fiber implanted over the SNr. **B.** Example behavioral session showing trial-by-trial licking activity (early lick trials excluded) for ipsilateral and contralateral optogenetic and baseline trials. Trials are arranged by optogenetic stimulation condition (stim trials top, baseline below). Blue and red horizontal lines depict the presentation of the air-puff during the sample period and yellow lines are the periods of optogenetic SNr inhibition. In this example, optogenetic inhibition of SNr shifts lick onset for contralateral trials closer to the start of the response window appears to slightly delay ipsilateral licking behavior. **C.** Average lick frequency traces for correct trials with (colored lines) and without (gray lines) unilateral optogenetic SNr inhibition. For contralateral trials (n = 71 opto, 140 baseline, **left**) and ipsilateral trials (n = 89 opto, 127 baseline, **right**) mice show faster decision licking activity on contralateral trials compared baseline lick decisions. Decision lick activity is depicted by the light shaded lines and retrieval licks by the dark shaded lines. **D.** Behavioral effects of unilateral SNr optogenetic inhibition in the withholding task. **E.** Changes in performance between contralateral (blue) and ipsilateral (red), off (light) and on (dark) stimulation. Bar height represents mean across all sessions (n = 16 sessions), with shapes representing the mean for each mouse (3 mice). Error bars represent SEM (bootstrap, 10000 iterations) and p values based on bootstrap (see methods, **p<0.01, *p<0.05). **Left.** Compared to baseline trials (light blue/red columns), licking performance during SNr inhibition (dark blue/red columns) produced a marginally significant increase in the percent of ipsilateral trials correct (p=0.0418, 16 sessions, 3 mice). **Middle Left.** Optogenetic inhibition of the SNr caused a significant increase in the percent of contralateral early lick fail trials (P=0.016) and a significant decrease in the percent of ipsilateral early lick trials (P<0.01). **Middle Right.** Optogenetic inhibition of the SNr increased the percentage of wrong direction ipsilateral trials (P<0.01). **Right.** SNr inhibition produced a slight trend and a marginally significant decrease percentage of no response trials for contralateral (P=0.23) and ipsilateral (P=0.048) trials, respectively. **H.** Mean reaction times for contralateral and ipsilateral trials with and without optogenetic SNr inhibition. Reaction times for contralateral trials were significantly decreased (P<0.01) and for ipsilateral trials were significantly increased (0.01) during SNr inhibition trials. **I.** Cumulative distribution plots showing the changes in the distribution of reaction times for contralateral (**left**) and ipsilateral (**right**) lick trials. Optogenetic stimulation trials are colored blue/red and baseline distributions are gray. The reaction time distribution shifts towards shorter reaction times for contralateral trials (**left**) and towards longer reaction times for ipsilateral trials (**right**).

### Unilateral excitation of the SNr impairs both contra- and ipsilateral licking activity

Since inhibiting basal ganglia output resulted in a contralateral bias in movement initiation and preparation, the classic model of basal ganglia rate coding leads to the prediction that increasing basal ganglia output would exert an opposite effect on licking activity, namely a reduction of correct decisions to lick in the direction contralateral to optogenetic stimulation. To test this prediction, we used the neural activator, ChR2(ET/TC), to increase output activity from the SNr (Fig 4A). The extent of the ChR2(ET/TC) opsin expression was again well localized to the SNr and SNr terminals observable in the VM thalamus and brainstem (typical example shown in Fig 4A, right). In awake mice, blue light (473 nm) stimulation of the SNr using a 1 s continuous pulse increased the already highly active SNr to approximately double the baseline firing rate for the duration of the light stimulation (mean firing rate at baseline = 13.9Hz vs. opto = 49.9Hz, n = 8 single units, Fig 4BC). In mice performing the anticipatory variant of the behavioral task (n = 4 mice, 16 sessions, Fig 4D-F), exciting the right SNr suppressed both contralateral and ipsilateral licking activity for the duration of the optogenetic stimulation after anticipatory licking had already started in the first 0.5 s of the air-puff (Fig 4EF). Following the offset of the optogenetic perturbation, mice often resumed licking towards the correct spout (Fig 4EF), indicating that they remembered the air-puff direction through the period of movement inhibition. However, successful task performance significantly decreased in both contralateral and ipsilateral lick trials with ChR2 stimulation (Fig 4H, P<0.01 (left), P<0.01 (right), bootstrap). This decrease in performance was both a result of licking the wrong spout direction (Fig 4H, P<0.01 (left), P<0.001 (right), bootstrap) and not licking during the response period (Fig 4H, P<0.05 (right), bootstrap), suggesting a partial loss of the neural representation of lick direction during the motor inhibition. In control mice injected with an EYFP virus without ChR2, blue light stimulation had no effect on licking behavior or task performance (n = 2, data not shown). These results give a more complex picture of bidirectional motor control with SNr inhibition or excitation than predicted by the classic rate model. Particular striking was a complete bilateral lick cessation during ChR2 induced SNr rate increases, which was distinctly different from the lateralized effects of ARCH inhibition.

**Figure 4.**
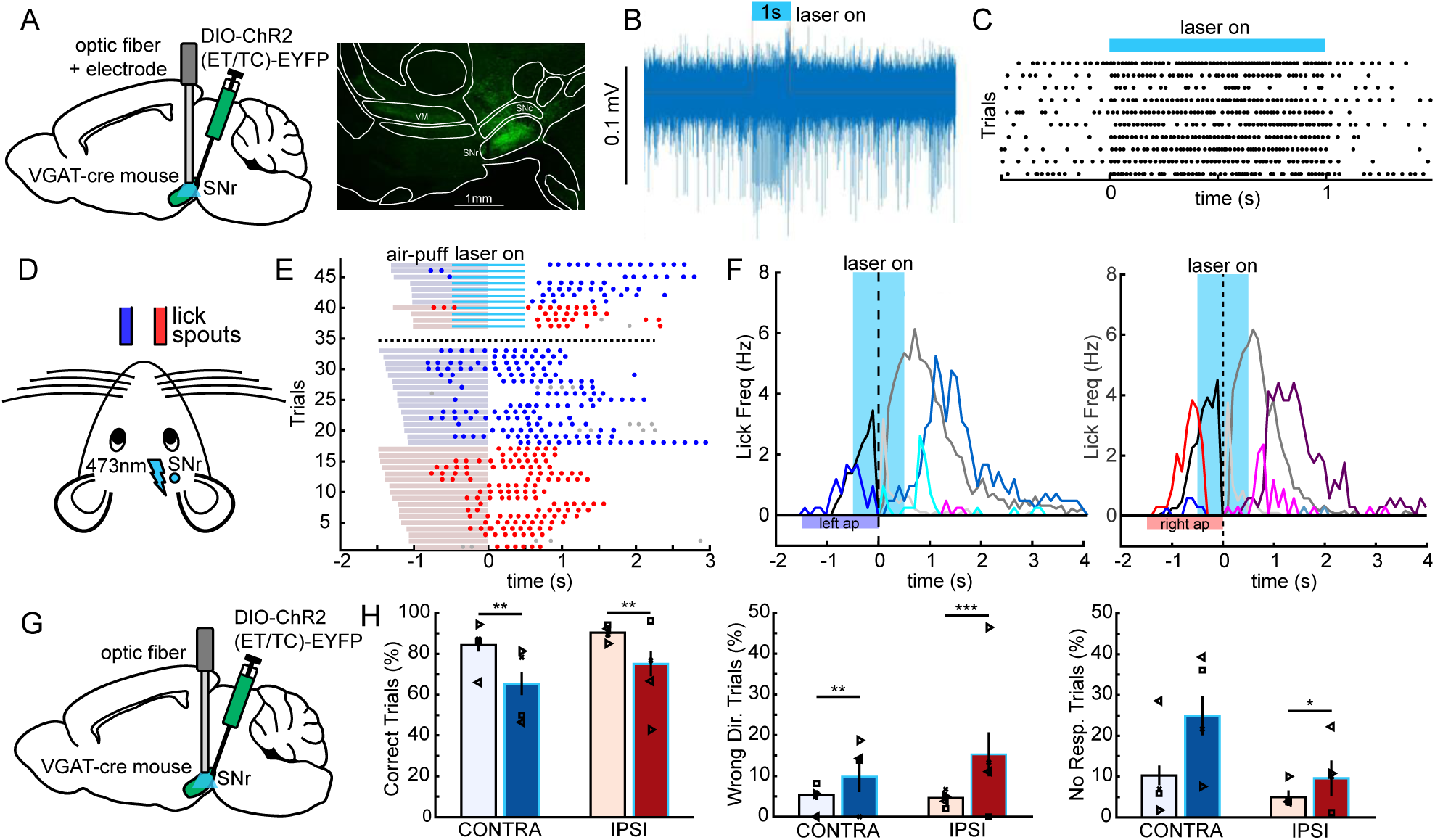
Unilateral excitation of the SNr impairs both contra- and ipsilateral licking activity. **A.** Diagram of optogenetic vector injection and optic fiber targeting. Cre-dependent AAV2-DIO-hChR2(E123T/T159C)-EYFP was injected in the right SNr of VGAT-cre transgenic mice and optic fiber attached to a microelectrode was lowered in to SNr. Example of ChR2(ET/TC)-EYFP expression (green) in SNr and axonal projections in VM thalamus and brainstem areas (right). **B.** Example multi-unit recording of an SNr neuron expressing ChR2(ET/TC). 473nm laser stimulation increases firing activity for the duration of the light pulse and firing quickly resumes following the offset of the light stimulation. **C.** 10 stimulation trials aligned to the onset of stimulation that show the excitation of SNr activity across trials (single unit isolated from recording shown in B). During ChR2 excitation, mean firing rate increased from 13.9Hz to 49.9Hz (n=10 neurons). Firing rate increase of example neuron is 14.4Hz vs. 46.3Hz during optogenetic stimulation. **D.** Illustration showing mouse orientation with respect to right and left lick spouts positioned in front of the mouse and optogenetic illumination through fiber implanted over the SNr. **E.** Example behavioral session showing licking activity for ipsilateral and contralateral optogenetic and baseline trials. Trials are arranged by optogenetic stimulation condition (stim trials top, baseline below) and length of air-puff presentation. Blue and red horizontal lines depict the presentation of the air-puff during the sample period and light blue lines are the periods of optogenetic SNr excitation. In this example, optogenetic excitation of SNr disrupts licking behavior both ipsilateral and contralateral to the side of optogenetic stimulation, though licking resumes following the offset of the light. **F.** Average lick frequency traces for correct trials with (colored lines) and without (gray lines) optogenetic SNr excitation. For contralateral trials (n = 96 opto, 168 baseline, **left**) and ipsilateral trials (n = 103 opto, 153 baseline, **right**) mice show anticipatory licking activity towards the correct spout that increases until the start of the response period for baseline licking, though is suppressed following the onset of light stimulation for optogenetic trials. Anticipatory licks are depicted by the middle colored line, followed by decision licks (light color) and retrieval licks (darkest color). **G.** Behavioral effects of unilateral SNr optogenetic excitation in the anticipatory task. **H.** Changes in performance between contralateral (blue) and ipsilateral (red), off (light) and on (dark) stimulation. Bar height represents mean across all sessions (n = 16 sessions), with shapes representing the mean for each mouse (4 mice). Error bars represent SEM (bootstrap, 10000 iterations) and p values based on bootstrap (see methods, ***p<0.001, **p<0.01, *p<0.05). **Left.** Compared to baseline trials (light blue/red columns), licking performance during right SNr excitation (dark blue/red columns) produced a significant decrease in the percent of contralateral and ipsilateral trials correct (p<0.01, 16 sessions, 4 mice). **Middle.** Optogenetic excitation of the SNr caused a significant increase in the percent of wrong direction fail trials for both contralateral (P<0.01) and ipsilateral (P<0.001) trials. **Right.** For contralateral lick trials, SNr excitation showed a trend towards increase percentage of no response trials (P=0.094) and a marginally significant increase in no response trial percentage for ipsilateral lick trials (P<0.014).

### Unilateral excitation of the SNr in ‘withholding’ task variant improves contralateral withholding of anticipatory licks and suppresses licking bilaterally

A subset of the ChR2 expressing mice (3 of 4) were further trained to perform the withholding version of the task. In agreement with the results for the anticipatory task, licking was generally suppressed until after the offset of the optogenetic stimulation (3 mice, 11 sessions, Fig 5BC. Unlike for the anticipatory variant of the task, however, SNr excitation led to an increase in percent correct for contralateral trials (Fig 5E, P<0.05). While this effect seems opposite to our hypothesis, examining the failure outcomes shows that this change was driven primarily by a decrease in the percent of contralateral early lick fails (Fig 5E, P<0.001), indicating an improvement in the ability to withhold premature licking predominantly to the contralateral side. Optogenetic excitation of the SNr also led to an increase in no response trials towards the direction contralateral to optogenetic stimulation (Fig 5E, P<0.05) and a trend towards decision licks in the wrong direction (Fig 5E, p=0.071). However, wrong decision licks were much reduced compared to the anticipatory variant of the task, suggesting that the additional training during the withholding stage had strengthened the neural representation of air-puff direction and made it more resilient to the ChR2 induced period of lick suppression. On trials where the mice did lick following the offset of the optogenetic stimulation, reaction time was significantly increased compared to control trials for both the ipsilateral and contralateral lick directions (Fig 5FG) due to the suppression of licking in the initial 0.5 s after air-puff offset when light stimulation was still on. Mice expressing EYFP, but no ChR2 in SNr did not demonstrate any changes in licking performance (n=2, data not shown). Overall, these results give further support to the strong bilateral motor suppression of unilateral SNr excitation.

**Figure 5.**
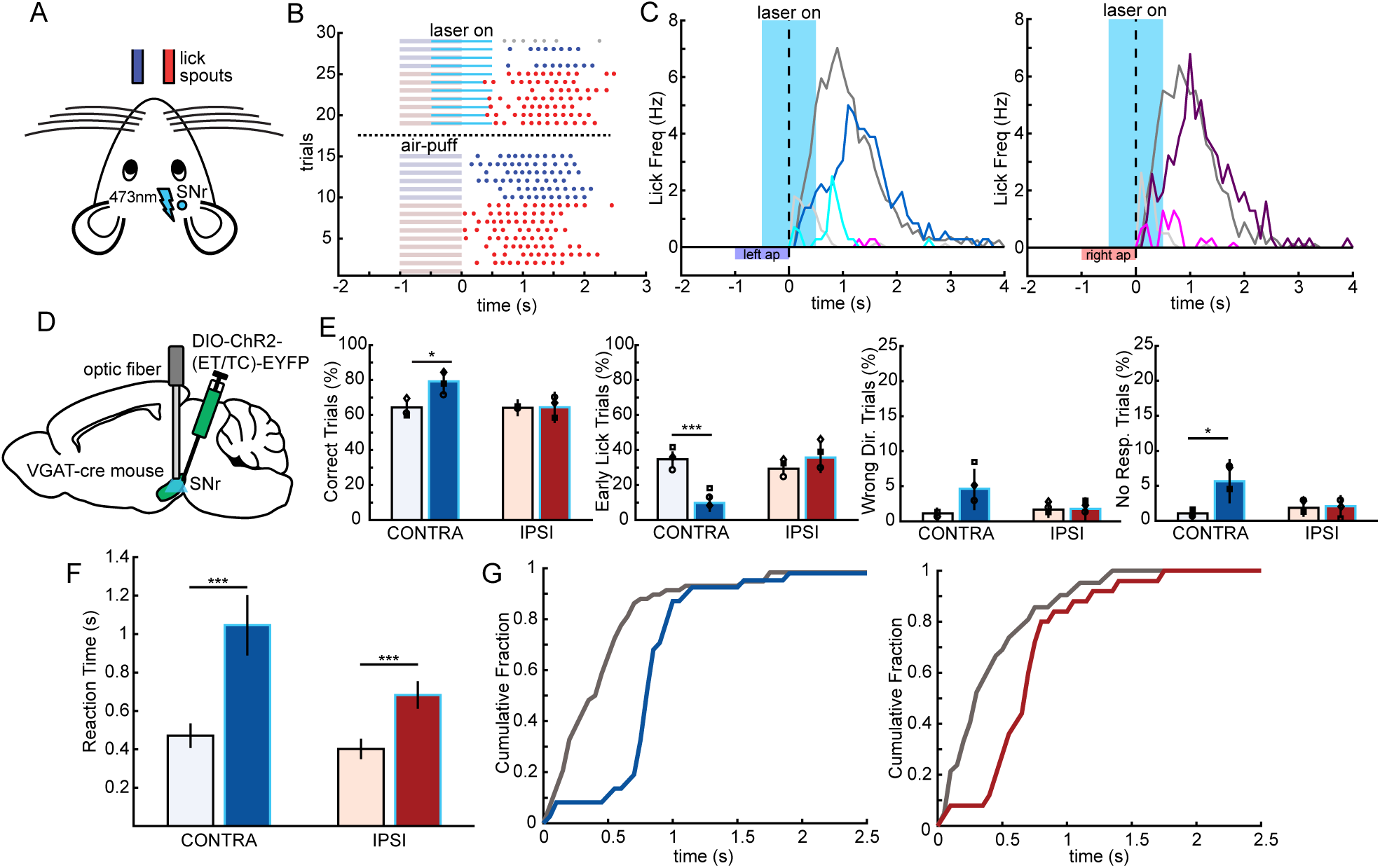
Unilateral excitation of the SNr increases reaction time towards both lick directions. **A.** Illustration showing mouse orientation with respect to right and left lick spouts positioned in front of the mouse and optogenetic illumination through fiber implanted over the right SNr. **B.** Example behavioral session during the withholding task showing trial-by-trial licking activity (early lick trials excluded) for ipsilateral and contralateral optogenetic and baseline trials. Trials are arranged by optogenetic stimulation condition (stim trials top, baseline below). Blue and red horizontal lines depict the presentation of the air-puff during the sample period and light blue lines are the periods of optogenetic SNr excitation. In this example, optogenetic excitation of right SNr delays lick onset for both ipsilateral and contralateral licking behavior. **C.** Average lick frequency traces for correct trials with (colored lines) and without (gray lines) optogenetic right SNr excitation. For contralateral trials (n = 63 opto, 132 baseline, **left**) and ipsilateral trials (n = 53 opto, 127 baseline, **right**) mice show delayed decision licking activity towards both spouts compared to baseline lick decisions. Decision lick activity is depicted by the light shaded lines and retrieval licks by the dark shaded lines. **D.** Behavioral effects of unilateral SNr optogenetic excitation in the withholding task. **E.** Changes in performance between contralateral (blue) and ipsilateral (red), off (light) and on (dark) stimulation. Bar height represents mean across all sessions (n = 12 sessions), with shapes representing the mean for each mouse (3 mice). Error bars represent SEM (bootstrap, 10000 iterations) and p values based on bootstrap (see methods, ***p<0.001, **p<0.01, *p<0.05). **Left.** Compared to baseline trials (light blue/red columns), licking performance during SNr excitation (dark blue/red columns) produced a significant decrease in the percent of contralateral and ipsilateral trials correct (p=0.023, 12 sessions, 3 mice). **Middle Left.** Unilateral optogenetic excitation of the SNr caused a significant decrease in the percent of contralateral early lick fail trials (P<0.001). **Middle Right.** Optogenetic excitation SNr did not produce any changes in direction licking behavior. **Right.** For contralateral lick trials, SNr excitation produced a significant increase percentage of no response trials (P=0.014). **H.** Mean reaction times for contralateral and ipsilateral trials with and without optogenetic SNr excitation. Reaction times for both contralateral and ipsilateral trials were significantly during SNr excitation (P<0.001). **I.** Cumulative distribution plots showing the changes in the distribution of reaction times for contralateral (**left**) and ipsilateral (**right**) lick trials. Optogenetic stimulation trials are colored blue/red and baseline distributions are gray. Both ipsilateral and contralateral lick responses are primarily delayed until after the offset of the optogenetic stimulation, shifting the distribution towards the ipsilateral spout compared to baseline responses.

### Unilateral excitation of nigrothalamic terminals in BGMT biases licking towards the ipsilateral direction in the anticipatory task variant

The differences in effects described for somatic SNr inhibition and excitation may be due to descending projections from the SNr towards the brainstem as well as ascending projections to BGMT. To begin to resolve the functional differences between these projection pathways, we selectively targeted the ascending pathway from the SNr to the BGMT. To achieve this, we stimulated ChR2 expressing SNr terminals in BGMT in the same mice used for somatic stimulation through a second fiber implanted over the BGMT. In contrast to somatic stimulation we found that SNr terminal stimulation in BGMT in the anticipatory variant of the task suppressed only contralateral licking behavior and did not affect ipsilateral licking (4 mice, 25 sessions, Fig 6BC). Successful trial execution was significantly impaired for contralateral lick trials but remained unchanged for ipsilateral licks (Fig 6E, P<0.001 (left), bootstrap). This change in performance was due to an increase of contralateral air-puff failed trials both by not responding (Fig 6E, P<0.001, bootstrap) as well as licking the wrong direction (Fig 6E, P<0.001, bootstrap). This last effect in particular was in stark contrast to the results with somatic SNr excitation, suggesting that the ascending thalamic pathway leads to a lateralized effect on motor preparation, whereas descending outputs may globally suppress the brainstem lick pattern generator.

**Figure 6.**
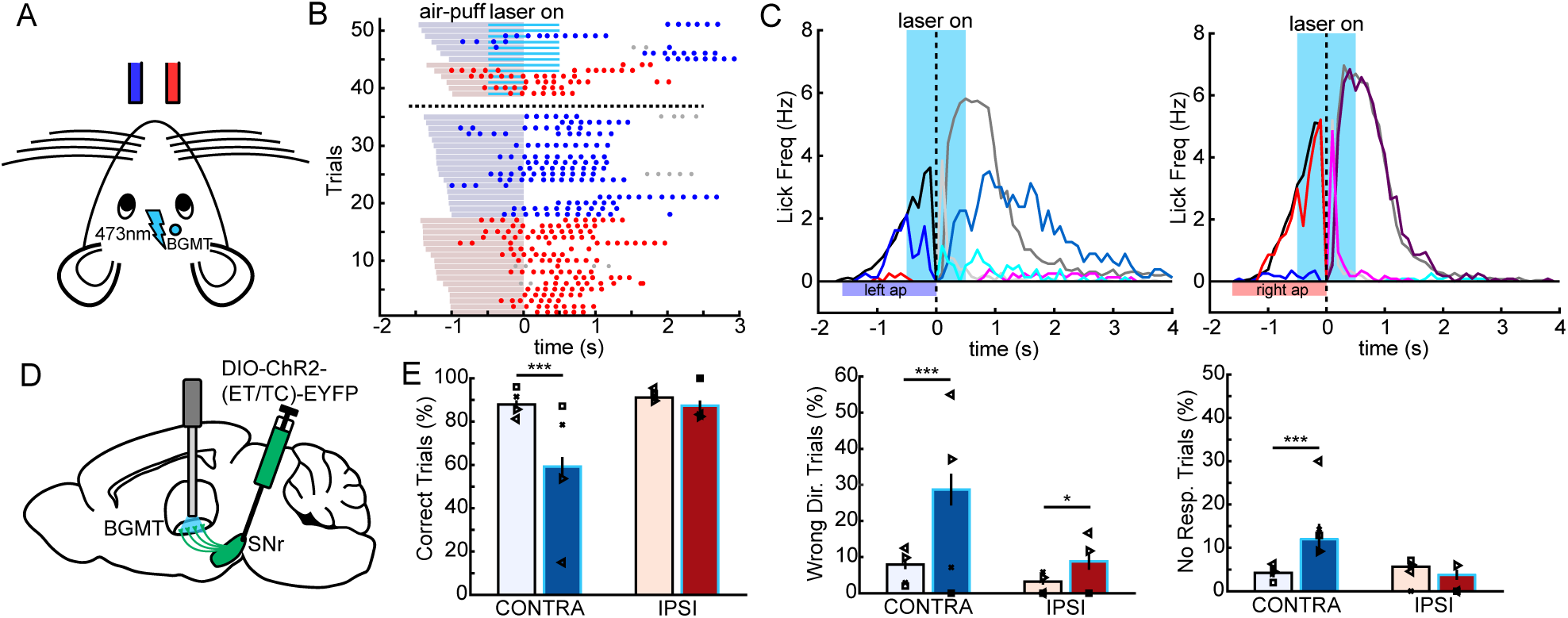
Unilateral excitation of SNr projections to the BGMT biases licking towards the ipsilateral direction. **A.** Diagram showing mouse orientation with respect to right and left lick spouts positioned in front of the mouse and optogenetic illumination through fiber implanted over the BGMT. **B.** Example behavioral session showing licking activity for ipsilateral and contralateral optogenetic and baseline trials during the anticipatory task. Trials are arranged by optogenetic stimulation condition (stim trials top, baseline below) and length of air-puff presentation. Blue and red horizontal lines depict the presentation of the air-puff during the sample period and blue lines are the periods of optogenetic SNr-BGMT excitation. In this example, optogenetic excitation of right SNr-BGMT projections disrupts only contralateral licking behavior during optogenetic stimulation trials. **C.** Average lick frequency traces for correct trials with (colored lines) and without (gray lines) optogenetic right SNr-BGMT terminal excitation. For contralateral trials (n = 95 opto, 310 baseline, **left**) and ipsilateral trials (n = 128 opto, 297 baseline, **right**) mice show anticipatory licking activity towards the correct spout that increases until the start of the response period for baseline licking, though anticipatory licking is suppressed following the onset of light stimulation for contralateral optogenetic trials. Anticipatory licks are depicted by the middle colored line, followed by decision licks (light color) and retrieval licks (dark color). **D.** Behavioral effects of unilateral SNr-BGMT optogenetic excitation in the anticipatory task. **E.** Changes in performance between contralateral (blue) and ipsilateral (red), off (light) and on (dark) stimulation. Bar height represents mean across all sessions (n = 25 sessions), with shapes representing the mean for each mouse (4 mice). Error bars represent SEM (bootstrap, 10000 iterations) and p values based on bootstrap (see methods, ***p<0.001). **Left.** Compared to baseline trials (light blue/red columns), licking performance during right SNr-BGMT terminal excitation (dark blue/red columns) produced a significant decrease in the percent of contralateral trials correct (p<0.001, 25 sessions, 4 mice). **Middle.** Optogenetic excitation of the right SNr-BGMT terminals caused a significant increase in the percent of contralateral (P<0.001) and ipsilateral (P=0.02) wrong direction fail trials. **Right.** For contralateral lick trials, SNr-BGMT terminal excitation caused a significant increase in the percentage of no response trials (P<0.001).

### Unilateral excitation of nigrothalamic terminals in BGMT in the withholding task variant facilitates suppression and increases reaction time for contralateral licking

Lastly, we again compared changes in the withholding task to those in the anticipatory variant. Licking activity was similarly impaired towards the contralateral direction while it remained intact towards the ipsilateral direction (3 mice, 13 sessions, Fig 7BC). The proportion of successful task performance in the ‘withholding’ variant was almost unchanged with SNr-BGMT terminal stimulation. However, there was a significant decrease in withholding failures (early lick trials) towards the contralateral direction (Fig 7E, P<0.05, bootstrap). This suggests SNr output to BGMT also contributes to the ability of the mouse to withhold movement, though only towards the contralateral direction. In this version of the task, lick reaction times in the response period to the contralateral direction were significantly longer than baseline (Fig 7FG, P<0.001, bootstrap). Ipsilateral reaction times were not significantly changed. This supports the conclusion that exciting SNr output specifically in BGMT exerted a lateralized influence on motor preparation and initiation.

**Figure 7.**
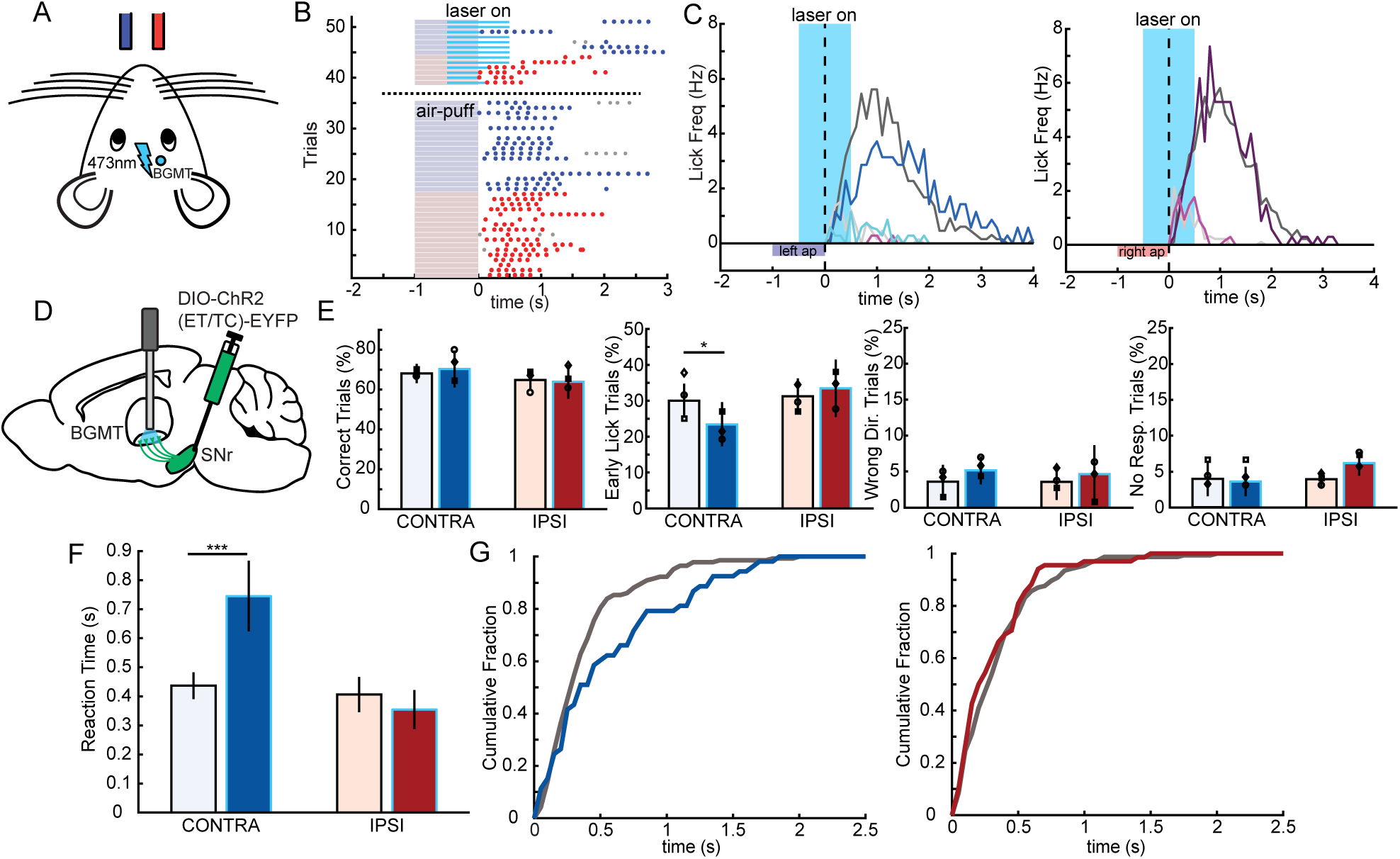
Unilateral excitation of nigrothalamic terminals facilitates suppression and increases reaction time for contralateral licking. **A.** Illustration showing mouse orientation with respect to right and left lick spouts positioned in front of the mouse and optogenetic illumination through fiber implanted over the SNr. **B.** Example behavioral session during the withholding task showing trial-by-trial licking activity (early lick trials excluded) for ipsilateral and contralateral optogenetic and baseline trials. Trials are arranged by optogenetic stimulation condition (stim trials top, baseline below). Blue and red horizontal lines depict the presentation of the air-puff during the sample period and light blue lines are the periods of optogenetic SNr-BGMT terminal excitation. In this example, optogenetic excitation of right SNr projections to BGMT delays lick onset of contralateral licking behavior. **C.** Average lick frequency traces for correct trials with (colored lines) and without (gray lines) optogenetic right SNr-BGMT terminal excitation. For contralateral trials (n = 87 opto, 245 baseline, **left**) and ipsilateral trials (n = 91 opto, 197 baseline, **right**) mice show similar licking activity during baseline trials, though during left optogenetic trials, decision and retrieval licking appears delayed. Decision lick activity is depicted by the light shaded lines and retrieval licks by the dark shaded lines. **D.** Behavioral effects of unilateral SNr-BGMT optogenetic terminal excitation in the withholding task. **E.** Changes in performance between contralateral (blue) and ipsilateral (red), off (light) and on (dark) stimulation. Bar height represents mean across all sessions (n = 13 sessions), with shapes representing the mean for each mouse (3 mice). Error bars represent SEM (bootstrap, 10000 iterations) and p values based on bootstrap (see methods, *p<0.05). **Left.** Compared to baseline trials (light blue/red columns), performance in optogenetic stimulation trials (dark blue/red columns) remains unchanged. **Middle Left.** Unilateral optogenetic excitation of the SNr projections to BGMT caused a significant decrease in the percent of contralateral early lick fail trials (P=0.023). **Middle Right.** Optogenetic excitation of SNr-BGMT did not produce any changes in percentage of wrong direction fail trials. **Right.** No response fail percentages were not altered by optogenetic stimulation of SNr projections to the BGMT. **H.** Mean reaction times for contralateral and ipsilateral trials with and without optogenetic SNr excitation. Reaction times for contralateral, but not ipsilateral trials were significantly increased during SNr excitation (P<0.001). **I.** Cumulative distribution plots showing the changes in the distribution of reaction times for contralateral (**left**) and ipsilateral (**right**) lick trials. Optogenetic stimulation trials are colored blue/red and baseline distributions are gray. Contralateral lick responses are delayed during optogenetic stimulation trials (**left**) while the response distribution for ipsilateral trials is unchanged compared to baseline (**right**).

## Discussion

Our results demonstrate that optogenetic excitation and inhibition of basal ganglia output from the SNr in mice drives lateralized and opposing influences on the control of directional licking. Specifically, unilateral inhibition of the SNr with ARCH activation leads to a decrease in performance of ipsilateral but an improvement in contralateral lick choices. The opposite performance effects were observed with activation of SNr terminals in BGMT (summarized in Fig. 8A,C). These findings indicate that the SNr firing rate decreases/increases are bidirectionally controlling lick direction choice. In mice performing a similar directional licking task, suppressing activity in either the VM thalamus, which is a major component of BGMT, or the ALM premotor cortex, through either optogenetic or muscimol inactivation selectively disrupts contralateral licking, while leaving ipsilateral licking unaffected (Li et al., 2015; Li et al., 2016; Chen et al., 2017; Guo et al., 2017; Svoboda and Li, 2017). Our results therefore suggest that this thalamo-cortical motor planning process can be gated by basal ganglia output in agreement with traditional rate coding concepts of basal ganglia – cortical loops (Alexander et al., 1990). According to this model, inhibiting the SNr with ARCH unilaterally leads to disinhibition of ipsilateral BGMT, and thus allows ipsilateral thalamocortical activity to develop and cause contralateral movement initiation. The opposite is expected for unilateral activation of the SNr with ChR2, and was observed in our study, but only if SNr terminals were activated in BGMT. Direct unilateral activation of GABAergic SNr cell bodies resulted in strong bilateral movement inhibition instead (Fig. 8B). A recent set of studies have implicated the SNr projection to the superior colliculus (the nigrotectal pathway) in the control of non-directional licking behavior (Rossi et al., 2016; Toda et al., 2017). In Rossi et al. 2016, researchers optogenetically bilaterally excited SNr projections at the level of the superior colliculus which led to diminished, though not completely suppressed, licking towards the lick spout positioned in front of the mouse. Our results showing that unilateral excitation of the SNr near completely suppressed licking could therefore be explained via projections to the SC. Consistent with this explanation, anatomical tracing experiments demonstrate that the SNr projects to SC bilaterally, while projections to the thalamus are ipsilateral (Deniau and Chevalier, 1992; Liu and Basso, 2008). Why somatic inhibition of the SNr did not facilitate licking bilaterally (opposite of SNr excitation) requires further examination but may suggest that the thalamic pathway is more susceptible to disinhibition than the nigro-collicular pathway.

**Figure 8.**
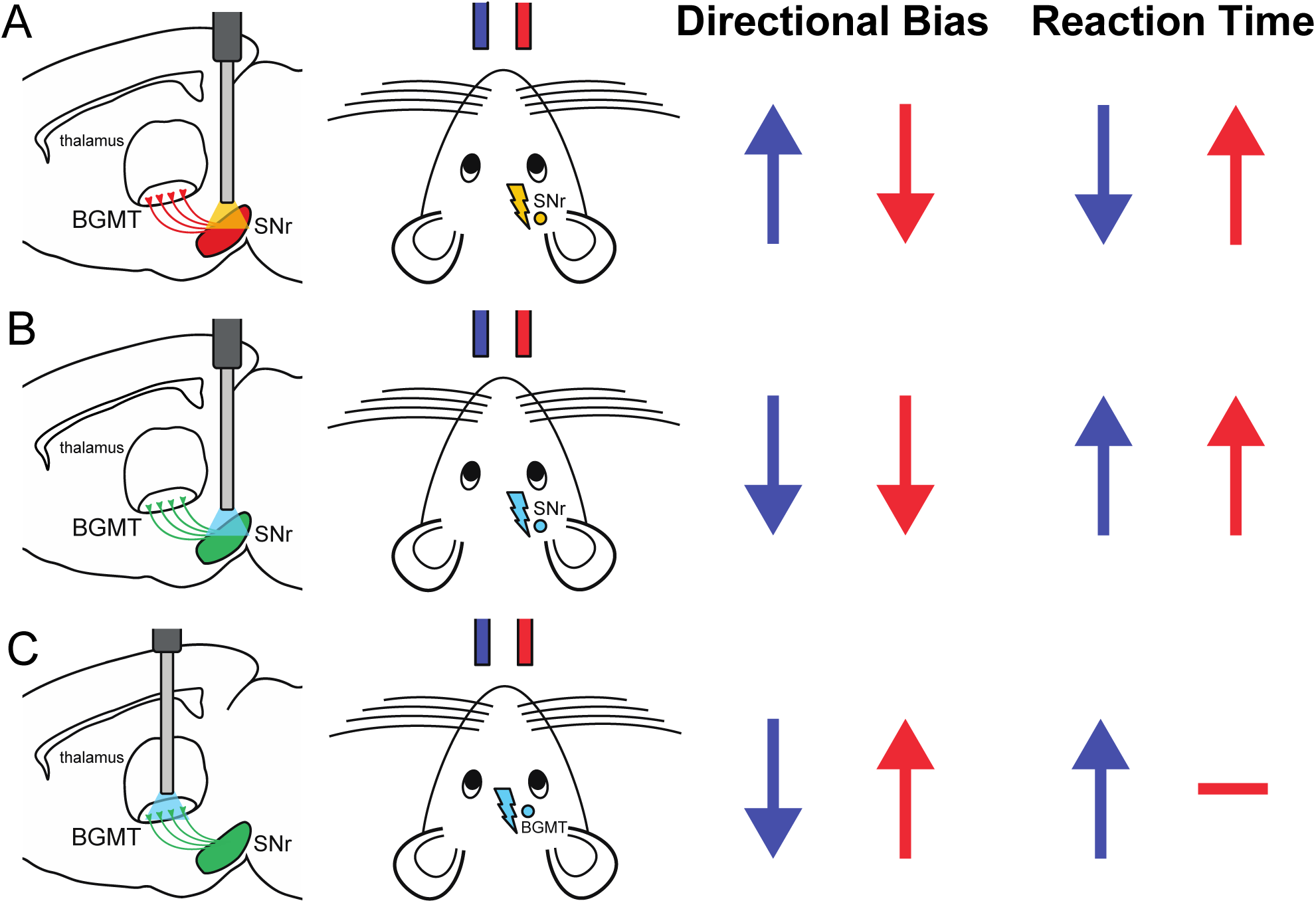
Summary of optogenetic manipulations and main results. **A.**Diagram depicting unilateral optogenetic inhibition of SNr expressing ARCH3 and orientation of lick spouts and mouse. SNr optogenetic inhibition is contralateral to the blue spout (left) and ipsilateral to the red spout (right). Optogenetic inhibition of the SNr produced a change in directional bias towards the contralateral spout (blue arrow) and away from the ipsilateral lick spout (red arrow). With respect to reaction time, optogenetic SNr inhibition reduced the reaction time towards the contralateral spout (blue) and increased time to respond towards the ipsilateral spout (red). **B.** Same as A., but for experiments with unilateral optogenetic excitation of SNr expressing ChR2(ET/TC). Optogenetic excitation of the SNr produced a bilateral change in directional bias; decreasing licking activity towards both the contralateral (blue) and ipsilateral (red) lick spouts. Additionally, optogenetic SNr excitation increased reaction times towards both contralateral and ipsilateral spouts. **C.** Same as A. and B., but for terminal excitation of SNr projections to the BGMT. Optogenetic excitation of SNr terminals in the BGMT produced a change in directional bias towards the ipsilateral spout (red) and away from the contralateral spout (blue). Excitation of SNr projections to BGMT also increased the reaction time towards the contralateral spout (blue) and did not affect reaction times towards the ipsilateral spout (red).

An important feature of our behavioral task training was the use of two different task variants that entailed different cognitive demands (Fig 1). The ‘anticipatory’ task mice were taught first, did not require lick withholding during air-puff delivery and indeed we observed a steady increase in anticipatory licking activity as the start of the response period was approached. This licking was in in the direction of the correct target (Fig 1C), therefore revealing the completion of air-puff stimulus evaluation and motor preparation as early as 1 s before air-puff offset (Fig 1C). Importantly, the prevalence of anticipatory licking was not affected by nigral ARCH activation (Fig. 2F), suggesting that tonic nigral inhibition in BGMT during reward anticipation was low when anticipatory licking was allowed, and therefore disinhibition had little effect on anticipatory licking. In contrast, a slowing of anticipatory licking was seen upon the onset of ChR activation of SNr terminals in BGMT for contralateral trials with a subsequent slowing of contralateral reward licks as well (Fig. 6C). This finding indicates that increased nigral activity in BGMT results in a lateralized slowing of movement. Several previous studies have shown that the basal ganglia exert lateralized control of motor circuits (Sakamoto and Hikosaka, 1989; Hikida et al., 2010; Tai et al., 2012; Dominguez-Vargas et al., 2017) and our results support this general concept. A slowing in licking is congruent with both the concepts of direct involvement of basal ganglia output in controlling movement velocity (Barter et al., 2015; Yttri and Dudman, 2016) or motor vigor (Dudman and Krakauer, 2016).

The ‘withholding’ variant of our lick task required mice to actively suppress licks for the duration of the air-puff (Fig 1D). By adding this withhold demand, we were able to investigate the role of the SNr in movement suppression and movement initiation by measuring early lick fail trials and reaction time following the offset of the air-puff, respectively. All mice preferred to lick in anticipation of directional choice behavior during initial training, and the suppression of this licking clearly required a cognitive effort. After the mice had learned this suppression, unilateral optogenetic inhibition of the right SNr suppressed anticipatory licking towards the ipsilateral lick spout, while it increased such licking to the contralateral spout (Fig. 3E). As described above, the same optogenetic inhibition of SNr activity in the same mice had not affected anticipatory licking in the ‘anticipatory’ variant of our task. In combination, these findings suggest that the mouse used increased nigral output to suppress anticipatory licking in the ‘withholding’ variant, and this increased nigral activity was inhibited with ARCH activation, which thus effectively undid the withhold learning. In addition, unilateral excitation of SNr terminals in the right BGMT now suppressed licking contralaterally (Fig 6E). These findings are in agreement with the active suppression of movement as a function of the indirect pathway in the basal ganglia (Nambu et al., 2002; Ozaki et al., 2017). Movement suppression is hypothesized to result from an activation of D2 receptor expressing GABAergic striatal projection neurons (SPNs), which then leads to increased activity in SNr and GPi via inhibition of the external pallidum (GPe) (Albin et al., 1989; Nambu et al., 2002). Cortical activity elicits excitation of GPi neurons more widely than inhibition, which is congruent with a center surround model of movement initiation and suppression (Ozaki et al., 2017). Optogenetic activation of D1 or D2 SPNs also supports the general principle that the indirect pathway causes movement suppression through SNr rate increases (Freeze et al., 2013), though optogenetic activation of either D1 or D2 SPNs resulted in excitation or inhibition of different SNr neurons.

Many studies have implicated the rodent SNr in the control of behavior other than licking, including for example locomotion (Roseberry et al., 2016) and head movement (Schmidt et al., 2013; Barter et al., 2015). To determine whether our manipulations inadvertently triggered such movements, we recorded high-speed video of the mouse pupil and face for a subset of trials across mice. Interestingly, we did not observe any movements (including eye, whisker, fore-limb) associated with our optogenetic manipulations (data not shown). This suggests that the behavioral impact seen with SNr optogenetic manipulations may be limited to behaviors trained in a reward-based task.

The BGMT with VM as its core component is well situated anatomically to influence ipsilateral cortical activity, with single-cell tracing studies in rodents depicting large projections branching across ipsilateral layer 1 of sensory and motor cortices (Kuramoto et al., 2009; Kuramoto et al., 2015). We showed here that activity in BGMT could be changed by inhibitory basal ganglia input and through this effect switch lateralized cortical decision-making processes about whisker stimulation dependent directional licking. Control of this behavior has previously been linked to a VM – ALM thalamocortical feedback loop (Li et al., 2015; Chen et al., 2017; Guo et al., 2017). Further, we also showed that lick movement vigor and reaction time can be impacted as well. The widespread axonal efferents of BGMT suggest that similar effects are likely for a multiplicity of other behaviors, though we saw no impact on spontaneous behavior. Because it is unlikely that a uniform optogenetic inhibition or excitation across SNr codes for any specific behavioral event, it seems most likely that our optogenetic stimulation interfered with endogenous population coding, which in a highly trained mouse will be quite specific to the trained behavior. However, even for certain behavioral events such as a stop-signal in a countermanding task, a relatively uniform activity increase has been observed across SNr neurons (Schmidt et al., 2013). The detailed mechanism by which SNr mediated activity changes in BGMT impact cortical decision making remains unclear, especially since layer 1 is far removed from the activity of cell bodies in L5/6 providing output from cortex. A likely candidate mechanism for amplifying such distal input is given by active distal dendritic properties such as NMDA spikes in L2/3 (Palmer et al., 2014) and calcium spike dependent BAC firing in L5 pyramidal neurons (Larkum et al., 1999, 2001; Larkum and Zhu, 2002). Similar active properties are also seen in the apical dendrites of L6 pyramidal neurons (Ledergerber and Larkum, 2010), which have a strong projection back to BGMT (Yamawaki and Shepherd, 2015). In addition, BGMT input could also act on L1 interneurons (Cruikshank et al., 2012), which provide a powerful inhibition of pyramidal neuron dendrites (Palmer et al., 2012; Palmer et al., 2013). The exact impact of BGMT input on these mechanisms and its integration into cortico-cortical information processing awaits future studies.

Author Contributions
A.M., P.B., C.W., G.S., and D.J. designed research; A.M., P.C., and C.B. trained mice and performed optogenetic experiments; A.M. analyzed data; A.M. and D.J. wrote the paper.

